# Detection of Plasmodium berghei infected Anopheles stephensi using near-infrared spectroscopy

**DOI:** 10.1101/195925

**Authors:** Pedro M. Esperança, Andrew M. Blagborough, Dari F. Da, Floyd E. Dowell, Thomas S. Churcher

## Abstract

**Background:** The proportion of mosquitoes infected with malaria is an important entomological metric used to assess the intensity of transmission and the impact of vector control interventions. Currently the prevalence of mosquitoes with salivary gland sporozoites is estimated by dissecting mosquitoes under a microscope or using molecular methods. These techniques are laborious, subjective, and require either expensive equipment or training. This study evaluates the potential of near infra-red spectroscopy (NIRS) to identify laboratory reared mosquitoes infected with rodent malaria.

**Methods:** *Anopheles stephensi* mosquitoes were reared in the laboratory and fed on *Plasmodium berghei* infected blood. After 12 and 21 days post-feeding mosquitoes were killed, scanned and analysed using NIRS and immediately dissected by microscopy to determine the number of oocysts on the midgut wall or sporozoites in the salivary glands. A predictive classification model was used to determine parasite prevalence and intensity status from spectra.

**Results:** The predictive model correctly classifies infectious and uninfectious mosquitoes with an overall accuracy of 72%. The false negative and false positive rates are, respectively, 30% and 26%. While NIRS was able to differentiate between uninfectious and highly infectious mosquitoes, differentiating between mid-range infectious groups was less accurate. Multiple scans of the same specimen, with repositioning the mosquito between scans, is shown to improve accuracy. On a smaller dataset NIRS was unable to predict whether mosquitoes harboured oocysts.

**Conclusions:** We provide the first evidence that NIRS can differentiate between infectious and uninfectious mosquitoes. Currently the method has moderate accuracy and distinguishing between different intensities of infection is challenging. The classification model provides a flexible framework and allows for different error rates to be optimised, enabling the sensitivity and specificity of the technique to be varied according to requirements.

## Background

The development and roll out of a simple to use rapid diagnostic test for human malaria has substantially improved monitoring of the disease [1]. There is an urgent need for a similar entomological tool to enhance mosquito surveillance and directly assess the impact of vector control interventions. The human force of infection is assessed using the entomological inoculation rate (EIR) which is calculated from the human biting rate and the proportion of mosquitoes with salivary gland sporozoites. Sporozoite prevalence is currently estimated either by manual dissection followed by visual observation using a microscope or through molecular methods such as *Plasmodium* specific PCR (polymerase chain reaction) or ELISA (enzyme-linked immunosorbent assay) which detect the circumsporozoite protein (CSP) [2, 3]. Dissection is laborious and requires staff with specialised training. Similarly, PCR requires well-equipped laboratories, expensive reagents, and technically trained staff. ELISA is more economic though it still needs expensive laboratory equipment and is thought to be unable to detect lightly infected mosquitoes [4]. The cost and laborious nature of these methods is compromising their systematic application as a large-scale monitoring tool.

Near-infrared spectroscopy (NIRS) is a fast, non-destructive and reagent-free scanning technique which has been shown to determine the age and species of morphologically indistinguishable mosquitoes of the *Anopheles Gambie s.l.* complex [5, 6], and to detect the presence of *Wolbachia* bacteria infections in *Aedes aegypti* mosquitoes [7]. The process involves scanning a mosquito at different wavelengths in the nearinfrared region of the electromagnetic spectrum to obtain their absorbance spectra. Differences in absorbance are indicative of differences in the molecular composition of the specimens scanned. Scans take a few seconds to be completed so that hundreds of mosquitoes can be scanned in the field each day by a single person without the need of a laboratory or extensive training. Following scanning, a calibration dataset is used to develop a predictive model to convert spectra into estimates of the characteristic under study (age, species or bacterial infection). Informative components of the spectrum are identified and used to predict the characteristic from an unknown sample.

Sporozoites are the most epidemiologically important *Plasmodium* life-stage, though there is utility in detecting other stages of the life-cycle. In the field, sporo-zoite prevalence is typically very low with ∼ 0–5% of caught mosquitoes being infectious [8]. This low prevalence means that many mosquitoes must be scanned to accurately estimate the percentage with the parasite. Being able to detect earlier mosquito-based parasite life-cycle stages, such as the presence of oocysts on the anopheline midgut wall, would increase the prevalence of the parasite in wild mosquito populations, meaning that sample sizes could be lower. Laboratory experiments also typically assess oocyst (as opposed to sporozoite) prevalence as it reduces rearing time, is safer as mosquitoes are not infectious and because most mosquitoes with oocysts go on to develop sporozoites [9]. There is increasing interest in quantifying the number/density of parasites in a mosquito and not just whether it is infected or not. Evidence indicates that highly infected mosquitoes are more infectious [10], and that parasite intensity might influence the efficacy of transmission blocking and pre-erythrocytic vaccines [10, 11].

This study investigates the use of NIRS to detect the presence of rodent malaria parasites in the *Plasmodium berghei* – *Anopheles stephensi* model system. Statistical methods for NIRS analyses are used to convert spectral data into estimates of sporozoite and oocyst prevalence and intensity and to prevent model overfitting.

## Methods

### Rearing

Colony mosquitoes were infected with rodent malaria as described previously [12]. Briefly, female mice (6–8 weeks old, Harlan, UK) were treated with phenylhydrazine and infected with 106 *P. berghei* PbPfs25DR3 three days later [12]. Colony *An. stephensi* mosquitoes (line SD 500, previously starved for 24 hours) were fed on infected mice three days later and unfed mosquitoes removed. After 12 days, a subsample of mosquitoes were killed using chloroform and kept on ice. Following killing, mosquitoes were scanned one by one and dissected using a light microscope to determine the number of oocysts on the midgut wall. All remaining mosquitoes were killed using chloroform 21 days post-feeding, scanned and the number of sporozoites in the salivary glands categorised on a logarithmic scale: 0 (no sporozoites), 1 (1–10), 2 (11–100), 3 (101–1000), 4 (>1000) [10, 13, 14]. All oocyst data were collected from a single feed on one cohort of mosquitoes whilst four replicates were used to generate the sporozoite data. The number of uninfected mosquitoes was augmented by adding mosquitoes fed on blood without the parasite, though were identical in every other way (same mosquito age, colony, blood-source).

### Scanning

Mosquitoes were scanned using a LabSpec4 Standard-Res *i* (standard resolution, integrated light source) near-infrared spectrometer and a bifurcated reflectance probe mounted 2mm from a spectralon white reference panel (ASD Inc. [15]). The machine records absorbance at 2151 wavelengths in the interval [350,2500] nanometers of the electromagnetic spectrum. All specimens were laid on their side under the focus of the light probe and spectra were recorded with RS3 spectral acquisition software (ASD Inc. [15]) which automatically records the average spectra from 20 scans. After each scan, mosquitoes were turned over onto their opposite side and rescanned to investigate whether multiple independent positioning and scanning improved overall accuracy. The light probe was centred on the head and thorax region of the mosquito though those scanned 12 days post-feed were also scanned centring on the abdomen region to investigate whether this part of the insect was more informative of oocyst load.

### Data analysis

A statistical machine learning approach is used to fit and cross-validate the best model using a generalised linear model (GLM) framework. A binomial logistic classification model is used to determine presence/absence of the parasite (two response classes: *y* = 1 for infectious/infected and *y* = 0 for uninfectious/uninfected) whilst a multinomial logistic classification model is used to investigate sporozoite intensity (which contains five response classes: *y* = {0,1, 2, 3, 4} for uninfectious, low, medium, high and very highly infectious).

Given that near-infrared (NIR) spectra are high-dimensional compared to the number of mosquitoes, we use partial least squares (PLS) to achieve dimension reduction and feature selection simultaneously [16]. PLS derives components which are used to transform the original spectra into PLS scores. The scores correspond to the reduced data and are used as covariates in the GLM to obtain the model parameter estimates. The number of PLS components used is a tuning parameter and is chosen by cross-validation to maximise predictive performance, as detailed below. To predict the infection status of a new sample from its spectra, we compute the linear predictor for that sample—which combines its spectra with the estimated GLM coefficients—and compare it to a threshold value. In the binomial case, the optimal threshold *t*_*_ is chosen to minimise the misclassification rate, such that the predicted class is then *infectious* if the linear predictor is larger than *t*_*_ and *uninfectious* otherwise; this is equivalent to obtaining the predicted class probabilities via the logistic transformation and determining an optimal threshold for this probability, although the first approach works better visually (shown in Figure 2C). In the multinomial case, the predicted class is simply equal to the class with the highest predicted probability.

**Figure 1:**
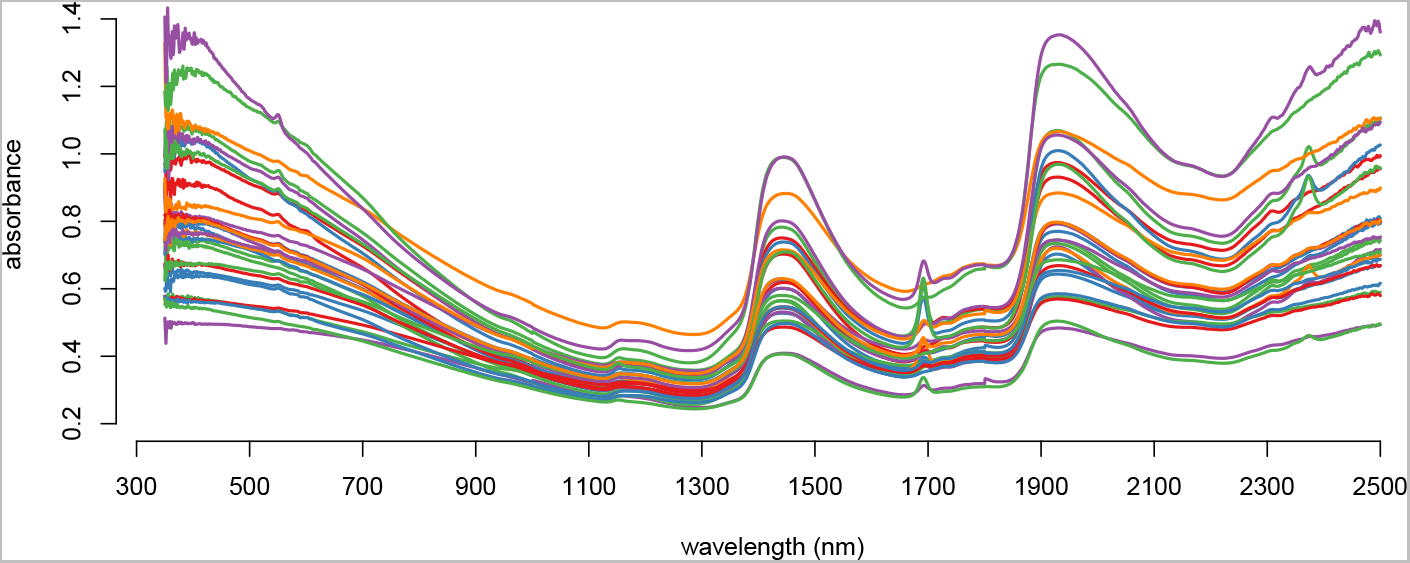
An illustration of NIR mosquito spectra. The colours of the 30 spectra denote the salivary gland sporozoite infection intensity on a log scale: 0 (orange), 1–10 (purple), 11–100 (green), 101–1000 (blue), >1000 (red).

**Figure 2:**
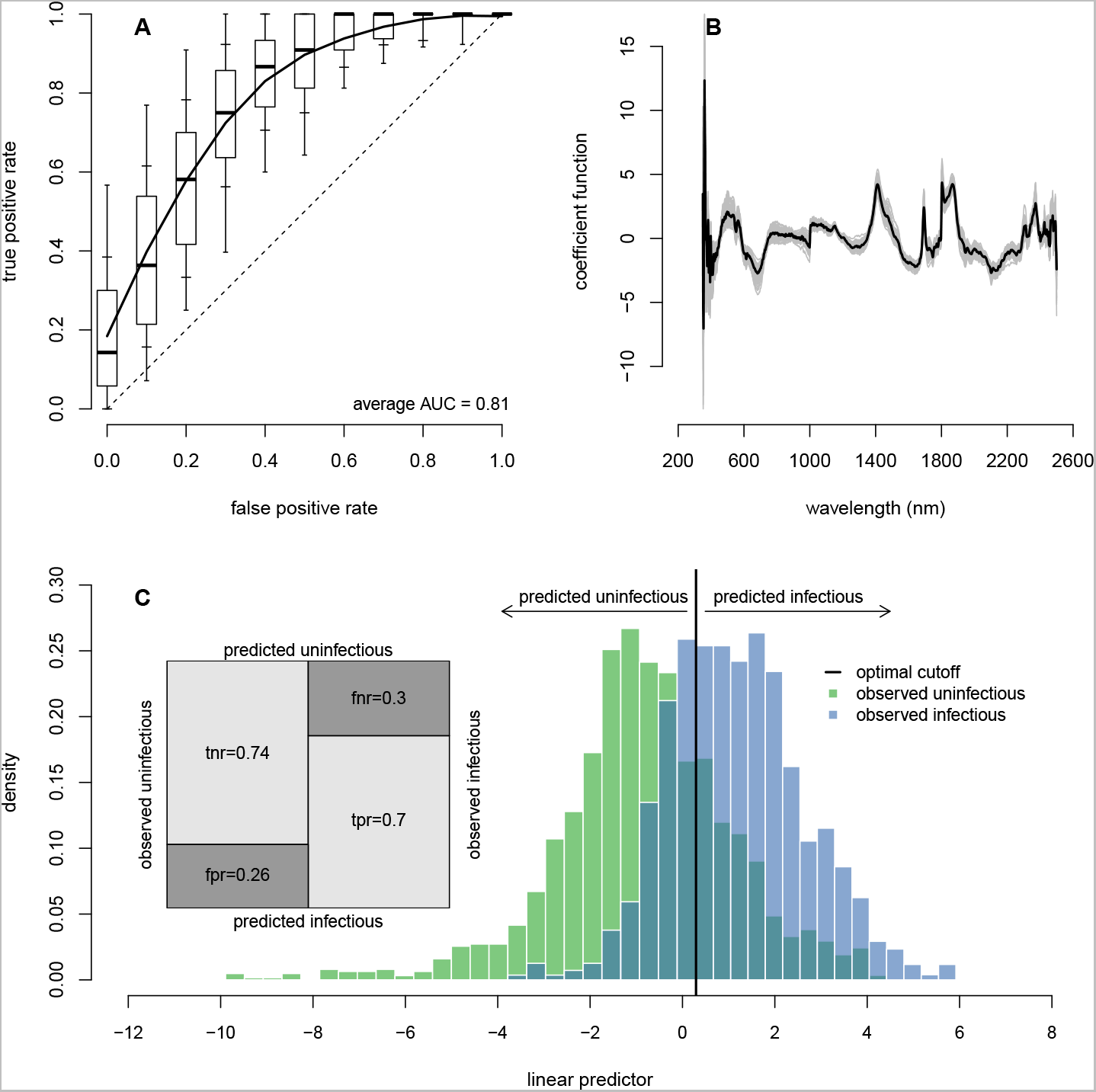
The ability of NIRS to determine sporozoite prevalence. Results are for a binomial GLM with 8 PLS components. [A] The receiver operating characteristic (ROC) curve for the best fit model showing the false positive and true positive rates achievable for different classification probability thresholds whilst the overall performance is given by the area under the ROC curve (AUC). The dashed line denotes a model with no predictive ability (a random chance of correctly predicting sporozoite presence) whilst a perfect model with 100% sensitivity and specificity would be in the top left corner (coordinates 0,1). The solid line shows the average ROC curve whilst the boxplots show the variability for 100 randomisations of the training, validation and testing datasets (with box edges, inner and outer whiskers showing 25th/75th, 15th/85th and 5th/95th percentiles, respectively; and the black line inside the box showing the median/50th-percentile). [B] The best fit coefficient functions for each of the 100 dataset randomisations (grey lines) and the corresponding average (black line). [C] The histogram of the estimated linear predictor for the test observations, colour-coded by the true class, shows the model’s ability to separate the two infection groups. The vertical black line indicates the optimum threshold for classifying mosquitoes as infectious or not. The shaded area where the two distributions overlap corresponds to misclassified test observations—false negatives to the left and false positives to the right of the optimal classification threshold. The confusion matrix (inset) shows the different error rates: `tnr` = true negative rate, `fnr` = false negative rate (specificity), `fpr` = false positive rate and `tpr` = true positive rate (sensitivity).

The area under the receiver operating characteristic (ROC) curve (AUC) is used to assess model accuracy and predictive performance, with a value closer to 1 indicating better performance. It gives the ability of a predictive model to correctly predict the true positive rate (sensitivity) and the true negative rate (specificity). In the multinomial case, when investigating sporozoite intensity, the AUC is computed by averaging the AUCs of all possible one-versus-all classification models, that is: dichotomising the response class into *y* = *c* versus *y* ≠ *c* and computing the standard two-class AUC, repeating the process for each class c, and averaging the results [17]. Misclassification rates are computed as the proportion of test observations incorrectly classified, given the optimal classification threshold that assigns equal weight to false negatives and false positives.

We use the standard three-step approach to build and asses the quality of a predictive model: training, validation and testing. Accordingly, the dataset is split into three subsets, each used at a different stage: (i) the training set is used to train a model with a given number of PLS components *K*, the procedure being repeated for different values of *K*; (ii) the validation set is used to evaluate each trained model to choose the optimal number of components (*K*_*_) which maximises the AUC; (iii) the testing set is used to evaluate the final model with *K*_*_ components in order to obtain an estimate of the generalisation error—an unbiased estimate of the error rate when the final model is used to predict a new (independent and identically distributed) observation. It is important that the final model is tested using data not previously used in either training or validation to avoid overfitting which inevitably leads to poor predictive (out-of-sample) performance [18].

The cross-validation results were averaged over 100 randomisations of the training, validation and testing datasets in order to average out sampling error. The optimal threshold for classification was chosen so as to minimise the error rate, giving equal weight to false positives and false negatives. The proportions of observations used in each subset were: 60% for training, 20% for validation, and 20% for testing. When more scans were conducted on an individual mosquito than were required for the analysis, scans were chosen at random.

The analysis was done using the `R` programming language [19] and the following packages: `glmnet` for the statistical models [20]; `ROCR` (for two-class problems) and `pROC` (for multi-class problems) to produce the ROC curves [21, 22]; and `pls` to derive the partial least squares components [23].

## Results

A total of 300 potentially infectious *An. stephensi* female mosquitoes were scanned, 138 (46%) of which had salivary gland sporozoites. A further 79 mosquitoes were scanned for oocyst detection, out of which 50 (63%) were confirmed to be infected with oocysts upon dissection. The sample sizes by infection levels and numbers of replicate scans are summarised in Table 1 and a sample of 30 spectra is depicted in Figure 1.

**Table 1:**
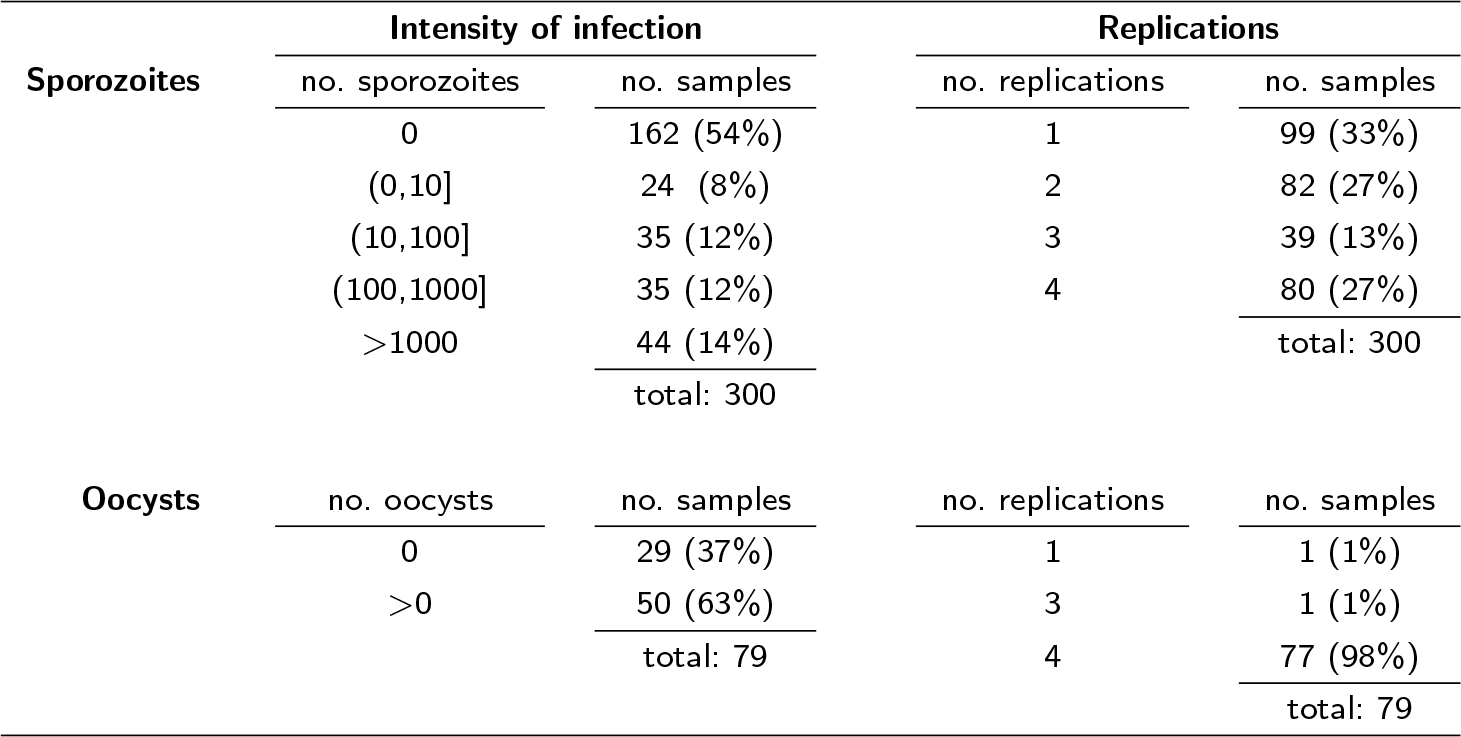
Sample sizes and replicate scans. Summary of the number of spectra collected by experiment (sporozoites or oocysts), the number of parasites in the mosquitoes (intensity of infection) and number of times the same mosquito was scanned in different positions (replications). Values in brackets indicate the percentage of the total samples with that intensity of infection or sample repetition.

Some specimens were scanned multiple times—with repositioning between replications—and their spectra averaged. Below we will show that this improves classification accuracy. Unless otherwise stated, the results presented include averaging of replicate scans.

### Sporozoite prevalence

NIRS can differentiate between infectious and non-infectious mosquitoes with high accuracy. The best fit model had 8 PLS components and an AUC of 0.81 (Figure 2A). Information provided by different regions of the spectrum can be visualised in the coefficient function (Figure 2B), with wavelength regions having larger magnitude being more important for the classification model. The distribution of the linear predictors for the test observations (Figure 2C) depicts the model’s capacity to separate infectious from uninfectious samples. The average classification accuracy is 72%, well above the frequency of the predominant class (54%; see Table 1) (i.e., if spectra is predictive of sporozoite prevalence, the model’s average accuracy rate must exceed the frequency of the predominant class). A detailed analysis of the classification results for test observations shows that true positive (infected) and true negative (uninfected) rates are, respectively, 74% and 70%, implying false positive and false negative misclassification rates of 26% and 30%, respectively.

A number of peaks were identified in the spectra at wavelengths 550, 1690 and 2370, which are not typically seen when mosquitoes are scanned. These peaks are in the region of the spectra consistent with samples contaminated with chloroform. It is hypothesised that this may have resulted from chloroform condensing over time in the petri dish when it was placed on ice. These peaks were predominantly present in one of the replicates and removing this replicate from the analysis had minimal impact on the overall accuracy or result (reducing the AUC from 0.81 to 0.78) and therefore it is unlikely to be causing the observed result.

### Sporozoite intensity

NIRS is not able to distinguish between mosquitoes with different numbers of salivary gland sporozoites with high accuracy. The average AUC among all one-versus-all models is 0.69 with the best fit model having varying success in categorising different infection groups, with AUC ranging from 0.56 for lowly infected to 0.80 for uninfected mosquitoes (Figure 3A).

**Figure 3:**
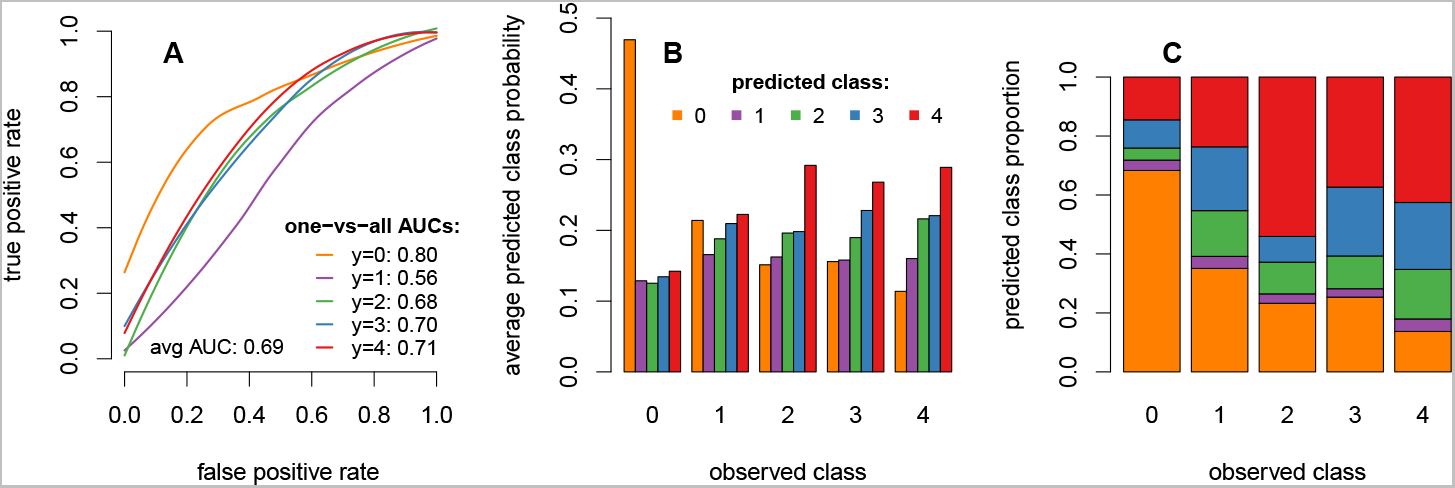
The inability of NIRS to determine sporozoite intensity. Results are for a multinomial GLM with 8 PLS components. [A] average receiver operating characteristic (ROC) curves for one-versus-all classification models and corresponding area under the ROC curve. [B] The average predicted class probabilities of test observations show that uninfectious mosquitoes are easily distinguished from infectious (large difference between the largest and second-largest average predicted class probabilities). Conversely, the model has difficulty in distinguishing between the different infection groups accurately (small differences between average predicted class probabilities). [C] The breakdown of predicted classes of test observations gives the matrix of misclassification rates. A class prediction corresponds to the class with the highest predicted probability so that panel C follows from panel B. The two panels give complementary information: whereas panel B hints at the difficulty in separating the different classes and can be used to pinpoint which classes are most confounded, panel C shows the actual proportions of misclassified test observation.

The model’s average predicted class probabilities for the extreme classes—uninfectious and very highly infectious—were relatively accurate (y = {0, 4}, respectively; Figure 3B). On average, the model estimates a high probability that a mosquito is uninfectious when it is in fact uninfectious. This probability decreases for more infectious mosquitoes as expected, that is, the more infectious a mosquito is, the lower is its estimated probability of being uninfectious. A similarly pattern can be observed for highly infectious mosquitoes.

However, distinguishing between the mid-range levels of infectiousness—low, medium and high—is less accurate (y = {1, 2, 3}, respectively; Figures 3B and 3C). The estimated probabilities for the low infection class (y=1) are uniform across all five levels of infection (difference between highest and lowest predicted probability equals 0.06), suggesting great uncertainty in predicting mosquitoes with low levels of infection (Figure 3B). The predicted classes for test observations suggest the same difficulty in predicting the middle infection groups, with misclassification rates between 77% and 96%. For uninfectious and very highly infectious mosquitoes, the misclassification rates are 32% and 57%, respectively (Figure 3C). Overall, the average misclassification rate is 53%, reflecting the difficulty in distinguishing mosquitoes with different infectiousness levels.

### Averaging specimens’ spectra

Scanning the same mosquito multiple times with repositioning of the mosquitoes before each replication improves the accuracy of the method. Table 1 indicates the number of repetitions for each experiment whilst Table 2 summarises the results of the effect on AUC and misclassification rate of averaging scans for each specimen. For sporozoites, averaging the spectra from two scans, chosen at random, for mosquitoes for which at least two scans were available (*n* = 201) substantially improved accuracy, with AUC increasing compared to single scans by 0.03 (a 4% improvement). Averaging all available scans for each mosquito (*n* = 300) resulted in an increase of 0.09 in the average AUC (a 12.5% improvement); and a decrease of 13 percentage points in the average misclassification rate.

**Table 2:**
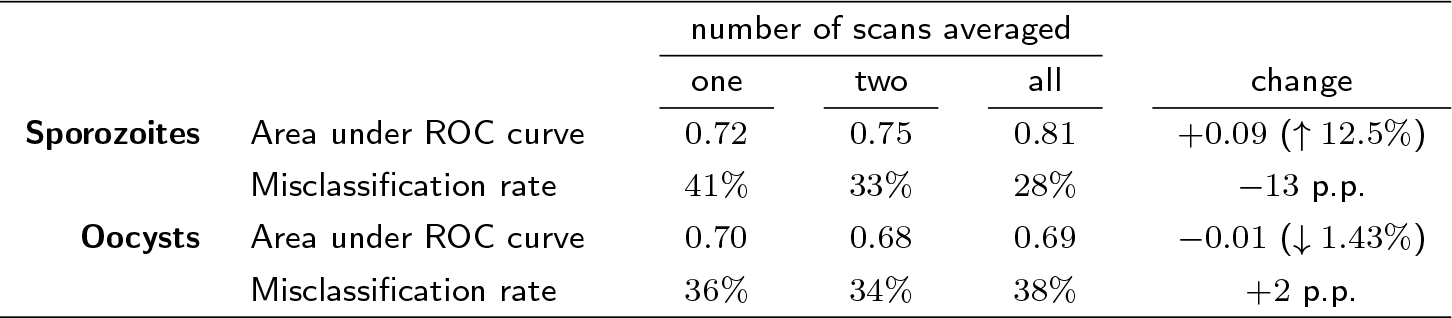
Improvements to the accuracy of NIRS in detecting presence of sporozoites and oocysts according to the number of times the mosquito was scanned. Mosquitoes are repositioned after each scan. AUC and misclassification rate when averaging one (no averaging), two and all available scans; see Table 1 for the number of replicates available. Improvements in performance, from one to all scans, are given in level for AUC and in percentage points for misclassification rate. Values in brackets indicate the percentage improvement in accuracy. Results are for a binomial GLM with 8 PLS components in both sporozoites and oocysts models.

|  |  | number of scans averaged |  |  | change |
| --- | --- | --- | --- | --- | --- |
|  |  | one | two | all |  |
| **Sporozoites** | Area under ROC curve | 0.72 | 0.75 | 0.81 | +0.09 (↑ 12.5%) |
|  | Misclassification rate | 41% | 33% | 28% | −13 p.p. |
| **Oocysts** | Area under ROC curve | 0.70 | 0.68 | 0.69 | −0.01 (↓ 1.43%) |
|  | Misclassification rate | 36% | 34% | 38% | +2 p.p. |

### Oocyst prevalence

The NIR spectra provided very little information on whether the mosquito was infected with oocysts. After averaging of mosquito spectra, the best model had an AUC of 0.69 and error rate very similar to oocyst prevalence; the average error rate was equal to 38% whereas the uninfected mosquitoes were 37% indicating little information in the spectra for predicting oocyst prevalence. Effectively, the best cross-validation model selects only two PLS components (the minimum possible value in our setup) which produces a very flat coefficient function, further supporting to this claim. Sample size was too low to investigate whether NIRS would be able to determine oocyst intensity.

For oocysts, averaging the spectra does not improve results, further supporting the previous results that the spectra is not predictive of oocyst load in this small sample (Table 2).

## Discussion

NIRS can differentiate between infectious and uninfectious mosquitoes with an overall accuracy of 72%. This is the first demonstration that NIRS can detect a malaria parasite in mosquitoes, but further work is needed to refine the technique before its measurement error and use can be fully understood. The accuracy is less than from the optimum recorded with PCR (the current accepted standard) though this technique has reproducibility issues in some laboratories and the practicalities and expense of this technique precludes its current widespread use in many settings ELISA is more economic though its sensitivity has been shown to perform with variable sensitivity in real world settings [24, 25].

The statistical machine learning approach to classification of NIR spectra provides a flexible framework and allows for different error rates to be optimised, enabling the sensitivity (true positive rate) and specificity (true negative rate) to be varied according to individual experimental requirements. In this study, we assigned equal weights to sensitivity (false positives) and specificity (false negatives), but one may wish to bias this criterion according to the question under investigation. For example, as an area nears malarial elimination, entomological surveillance becomes increasingly difficult as many thousands of mosquitoes need to be examined to accurately estimate the sporozoite rate. In this situation the classification threshold could be changed to minimise the false negative rate to ensure that all mosquitoes classified as negative were truly uninfectious. This would allow the proportion of mosquitoes with sporozoites to be estimated in a two-step process; 1) mosquitoes are quickly scanned to remove those known to be negative, 2) those remaining would be analysed by PCR to ensure all positive samples are identified. Such a method could substantially reduce entomological surveillance costs without compromising accuracy.

This work demonstrates the ability of NIRS to detect malaria in a laboratory rodent model system and the study needs to be repeated in natural parasite-vector combinations of medical importance. Nevertheless the *An. stephensi* – *P. berghei* system is widely used to understand the biology of the passage of the parasite through the mosquito and in the development of anti-malarial transmission-blocking drugs and vaccines so this work has direct biological relevance. These experiments require use of the standard membrane feeding assay where mosquitoes are fed on infectious blood before being individually dissected by hand under a microscope [26]. Dissection is slow, laborious, and inherently subjective so the use of NIRS may make it easier to screen large libraries of drug and vaccine candidates.

Distinguishing between different levels of infection in mosquitoes using NIRS is a challenging task. The differences in the spectra of these infection groups are subtle and there is considerable uncertainty in the classification of test observations (Figure 3). This uncertainty is likely to be due, at least in part, to the small sample sizes available (*n* = 300 for the sporozoite data). For instance, there are only 24 samples from the low infection group, corresponding to 8% of the total. Additionally, the dataset is highly unbalanced towards uninfected mosquitoes, which are the most accurately predicted of all infection groups (Table 1 and Figure 3). Larger sample sizes will be required to calibrate a more robust predictive model that can be used in real-life situations. A larger study will also be required to confirm that NIRS is unable to detect *P. berghei* oocysts as only 79 mosquitoes were available for analysis here.

The mechanisms by which NIRS detects the malaria parasite remain unknown and it is unclear whether the regions of the spectra identified as important are detecting parasite biomass, a mosquito response to the parasite, or a combination of both. Given the relative masses of the parasite to the insect it is likely that NIRS is detecting a mosquito response though further work is needed to clarify the mechanisms involved.

## Conclusions

We provide the first evidence that NIRS can be used to distinguish mosquitoes infectious with malaria from those which were not. The experiment must be repeated with wild natural parasite-vector combinations before its practical use as a tool for monitoring of vector control interventions can be assessed.

## Abbreviations

AUC: area under the (ROC) curve
GLM: generalised linear model
NIR: near-infrared
NIRS: near-infrared spectroscopy
PLS: partial least squares
ROC: receiver operating characteristic.

## Availability of data and materials

The datasets supporting the conclusions of this article are available in the Zenodo repository, https://doi.org/10.5281/zenodo.1001720. [27]

## Author’s contributions

Study conceptualization, funding acquisition and project supervision by AMB, DFD, TSC. Data collection by PME, AMB, TSC. Data analysis by PME. Writing by PME (original manuscript) and PME, AMB, DFD, FED, TSC (review and editing). All authors read and approved the final manuscript.

## Acknowledgements

The work was supported by UK Medical Research Council (MRC) Project Grant (MR/P01111X/1) and the MRC / UK Department for International Development (DFID) under the MRC/DFID Concordat agreement. The authors would like to thank Maggy T. Sikulu—Lord for technical advice. Mention of trade names or commercial products in this publication is solely for the purpose of providing specific information and does not imply recommendation or endorsement by the U.S. Department of Agriculture. USDA is an equal opportunity provider and employer.

